# Monet: An open-source Python package for analyzing and integrating scRNA-Seq data using PCA-based latent spaces

**DOI:** 10.1101/2020.06.08.140673

**Authors:** Florian Wagner

## Abstract

Single-cell RNA-Seq is a powerful technology that enables the transcriptomic profiling of the different cell populations that make up complex tissues. However, the noisy and high-dimensional nature of the generated data poses significant challenges for its analysis and integration. Here, I describe *Monet*, an open-source Python package designed to provide effective and computationally efficient solutions to some of the most common challenges encountered in scRNA-Seq data analysis, and to serve as a toolkit for scRNA-Seq method development. At its core, Monet implements algorithms to infer the dimensionality and construct a PCA-based latent space from a given dataset. This latent space, represented by a *MonetModel* object, then forms the basis for data analysis and integration. In addition to validating these core algorithms, I provide demonstrations of some more advanced analysis tasks currently supported, such as batch correction and label transfer, which are useful for analyzing multiple datasets from the same tissue. Monet is available at https://github.com/flo-compbio/monet. Ongoing work is focused on providing electronic notebooks with tutorials for individual analysis tasks, and on developing interoperability with other Python scRNA-Seq software. The author welcomes suggestions for future improvements.

## Introduction

Single-cell RNA-Seq (scRNA-Seq) has become a widely used technology to elucidate the transcriptomes of individual cell populations in complex tissues, with applications in immunology, cancer research, developmental biology, neurobiology, and other fields. The analysis of scRNA-Seq data presents a unique combination of computational and statistical challenges^1–4^: First, the data is very noisy, mostly due the fact that only a random subset of mRNA molecules from each cell is detected. Therefore, all scRNA-Seq analysis methods must adopt strategies aimed at separating biological expression differences from technical noise. Second, the data is very high-dimensional, making it essential to employ some form of dimensionality reduction. In high-dimensional space, cells all appear nearly equidistant from one another, an effect sometimes referred to as the ‘curse of dimensionality’. Third, datasets are very large, often containing data for thousands of cells, making it difficult to efficiently store and load data, both on-disk or in-memory. Fourth, in addition to the biological heterogeneity present within one dataset, researchers are commonly interested in studying heterogeneity across datasets (e.g., differences between treatment conditions or individuals), posing challenges as to how to jointly analyze, or “integrate”, multiple datasets. This requires methods that can overcome or correct for batch effects, the nature and magnitude of which are often unknown.

These technical challenges underlie and permeate almost any aspect of scRNA-Seq data analysis, independently of whether the ultimate goal is to obtain a particular visualization of the data, to perform clustering, to order cells along a developmental trajectory, or to make comparisons between datasets. Since there often exist many different approaches for each type of analysis (e.g., many different clustering algorithms), and many different approaches to address each of the aforementioned technical challenges, it is perhaps not surprising that hundreds of scRNA-Seq analysis tools have been developed^5^. However, even for experienced computational biologists, navigating this vast methodological landscape can be difficult, as it often requires significant effort to understand how the approaches chosen by a particular tool affect the data and interact with each other to produce the final analysis result.

To allow researchers to perform common scRNA-Seq analysis tasks without having to navigate hundreds of different tools, multiple “comprehensive” software packages for analyzing scRNA-Seq data have been developed. The most popular examples include the R packages Seurat^6^ and Monocle^7^, as well as the Python package Scanpy^8^. In principle, these packages can implement a “core analysis framework” for addressing the aforementioned technical challenges, while providing a user-friendly interface for performing different scRNA-Seq analysis tasks. To be able to properly interpret analysis results, researchers need to develop an understanding of how the core analysis framework operates, at least at an intuitive level. However, it is much more feasible to familiarize oneself with a single framework than with dozens of independently developed tools with narrower focus. Package authors should therefore publish clear explanations of the core analysis framework.

Here, I describe a new Python software package termed *Monet* for analyzing scRNA-Seq data. The core analysis framework of this package consists of an algorithm to learn a PCA-based latent space from a given dataset, with the dimensionality being automatically determined using molecular cross-validation^9^, as well as an algorithm to project arbitrary scRNA-Seq datasets (usually from the same tissue) into such a latent space. While PCA is commonly used in the analysis of scRNA-Seq data^10^, Monet’s core analysis framework avoids or replaces many of the steps commonly used by other packages in the preprocessing of the data, including gene selection, log transformation, or any kind of parametric modeling^2,11^. It also explicitly puts latent spaces, encapsulated by *MonetModel* objects, at the center of the analysis of scRNA-Seq data. Monet relies as much as possible on standard machine learning algorithms to perform specific tasks (e.g., visualization with t-SNE, clustering with DBSCAN, K-nearest-neighbor classification for label transfer), while also implementing successful ideas previously described in the single-cell literature (e.g., batch correction by matching mutual nearest neighbors^12^). The package also contains an implementation of *ENHANCE*, a previously developed denoising method^3^ that uses the Monet latent space model (with a simpler heuristic for inferring dimensionality) in its k-nearest neighbor aggregation step.

## Results

### Monet leverages Python tools for data manipulation, machine learning and visualization

To develop a software for analyzing scRNA-Seq data in Python, I relied on successful open-source packages from the Python ecosystem (**Figure 1a**). Expression matrices in Monet are represented using the *ExpMatrix* class, which is a subclass of the pandas *DataFrame* class. To store and load raw scRNA-Seq data consisting of UMI counts for each gene and each cell, I found that *numpy*’s compressed.*npz* binary format offers much better performance in terms of disk usage and loading times than plain-text formats. The *save_npz()* and *load_npz()* functions of ExpMatrix objects use this format to efficiently save and load data to/from the hard drive, respectively. For statistical and machine learning tasks, Monet relies heavily on *scikit-learn* and *scipy*, while *plotly* is used to generate visualizations that can be embedded into *Jupyter* notebooks. These packages offer an incredibly broad set of features and are actively maintained and developed. They also can be easily installed using the *conda* package manager, although Monet currently only supports installation with the *pip* package manager. Work to make Monet installable with conda is ongoing.

**Figure 1:**
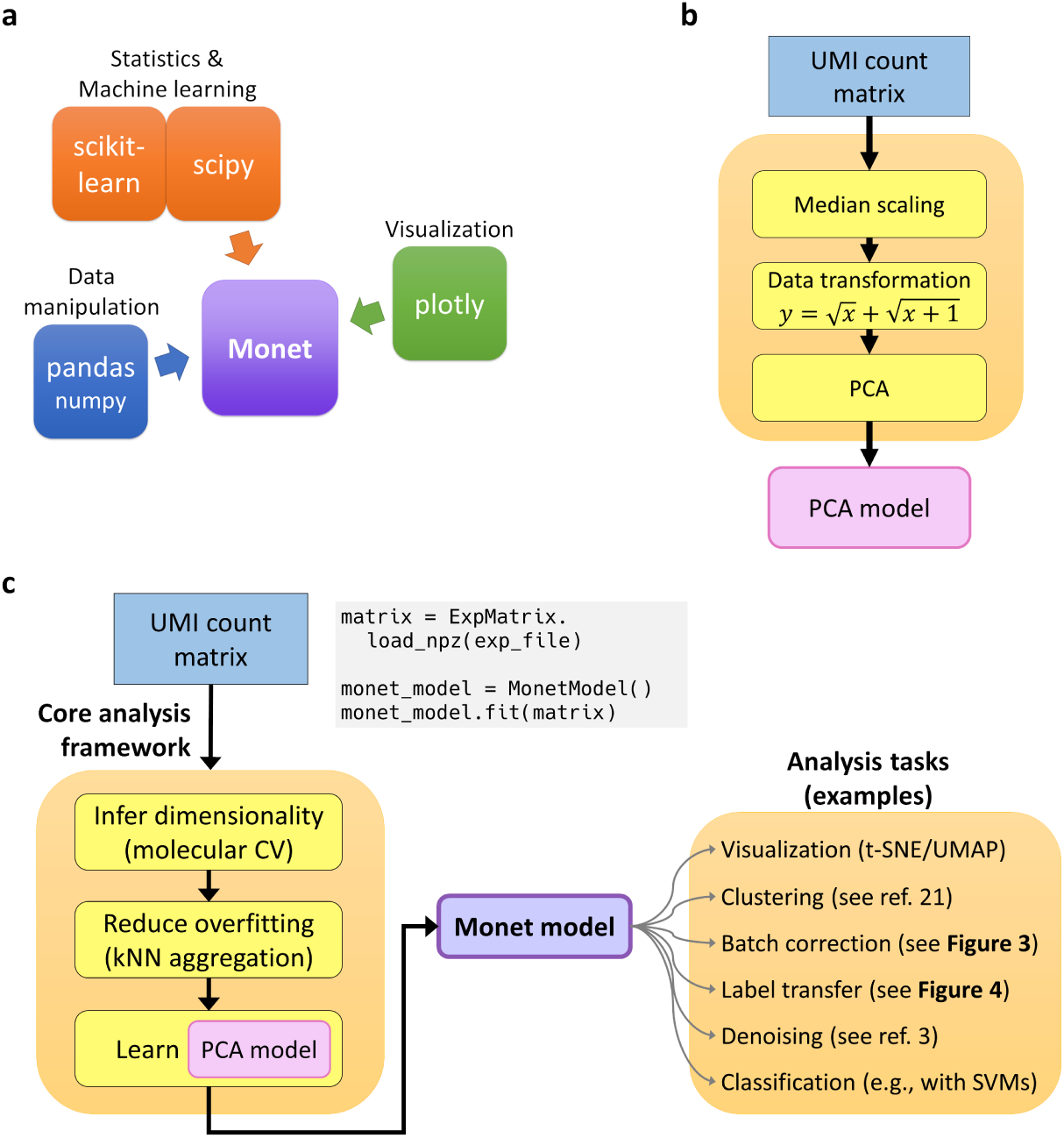
Design elements of the Monet package for scRNA-Seq data analysis. **a** Depiction of third-party Python packages used to implement key functionalities. **b** The core analysis framework relies on the application of PCA to the UMI count matrix, after applying median scaling a simple square root-based data transformation. **c** Overview of the core analysis framework, potential analysis tasks, and code examples. A Monet model is obtained by inferring the dimensionality using molecular cross-validation, applying k-nearest neighbor aggregation, and then performing PCA on the aggregated (and re-scaled) data. This model then serves as the basis for various downstream analysis tasks.

### Monet’s core analysis framework relies on simple data transformations and PCA

Gene expression measurements obtained from scRNA-Seq, represented by UMI counts, are associated with significant levels of technical noise. The amount of noise strongly depends on the expression level of the gene, but in many cases exceeds 100% (coefficient of variation), in which case the standard deviation representing the technical variation is larger than the true expression level. In 2014, Grün et al.^1^ observed that the technical noise displayed by UMI counts can be understood as a combination of *sampling noise* and *efficiency noise*, where sampling noise refers to the stochastic variation introduced because only a small random subset of transcripts for each cell is detected, whereas efficiency noise refers to stochastic differences in the overall number of transcripts detected for each cell. The authors further observed that sampling noise was the dominant source of technical variation for all except the most highly expressed genes. In a 2017 paper^13^, I built on these observations and proposed to preprocess scRNA-Seq datasets by using a two-step procedure (**Figure 1b**). In the first step, the expression profiles of all cells are scaled to the median transcript count per cell, in order to counteract efficiency noise, which was already discussed by Grün et al. In the second step, a simple square root-based transform, 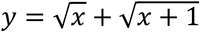, is applied to the scaled expression values. The main motivation for using this *Freeman-Tukey transform*^14^ is to let measurements contribute to the downstream PCA step in approximate proportion to their signal-to-noise ratio, meaning that the relatively accurate measurements obtained for highly expressed genes contribute more (but not too much) to the analysis than those of lowly expressed genes, which contain very little information. This simple transform therefore obviates the need for a gene selection step, which is used by many scRNA-Seq analysis tools^2^. The reader may refer to the **Methods** section for a discussion of the effects of different data transformations. After scaling and applying the FT transform, Monet performs principal component analysis (PCA) on the data, using a fast randomized implementation provided by scikit-learn based on algorithms described by Halko et al.^15^ Monet complements this simple approach to performing PCA on scRNA-Seq data with algorithms for inferring the dimensionality of a dataset and for reducing PCA overfitting, which are described below. The resulting Monet model represents a tissue-specific latent space that can form the basis for many different analysis tasks (**Figure 1c**).

### Monet infers the dimensionality of a dataset using molecular cross-validation

Methods aimed at constructing a latent space for scRNA-Seq data are faced with the challenge of how to appropriately infer the number of dimensions to use. Recently, Batson et al.^9^ proposed an approach they termed *molecular cross-validation* (MCV) that enables the systematic estimation of model parameters in an unsupervised setting. The idea behind MCV is to split the dataset into a training and a test set, by partitioning the individual mRNA molecules observed for each cell. The authors used both simulation studies and statistical theory to show that when this is done in an appropriate fashion, the parameter value that minimizes a loss function on the test dataset (MCV loss) is also the value that minimizes the ground truth loss. Monet implements 5-fold MCV with a Poisson loss function to infer the dimensionality in the context of the previously described PCA framework (see **Methods**).

I first tested this approach on three different human PBMC datasets obtained using 10x Genomics’ Chromium technology (**Figure 2a**). For the two PBMC datasets obtained using the v2 chemistry, Monet inferred a significantly lower dimensionality (19 and 22) than for the dataset obtained using the v3 chemistry (30). This is consistent with the fact that the v3 chemistry achieves a much higher transcript detection rate, and therefore is able to produce a higher-resolution view of the different subpopulations of cells. Both v2 PBMC datasets were obtained using cells from the same donor, and only differed in the number of cells profiled. Monet inferred a slightly lower dimensionality (19) for the dataset containing ∼4,000 cells than for the dataset containing ∼8,000 cells (22), which again seemed consistent. Finally, I tested the approach on a mouse embryonic heart dataset, for which Monet inferred a dimensionality of 52. This was significant higher than for any of the PBMC datasets, which was consistent with the fact that the heart dataset appeared to contain a much larger number of distinct cell types.

**Figure 2:**
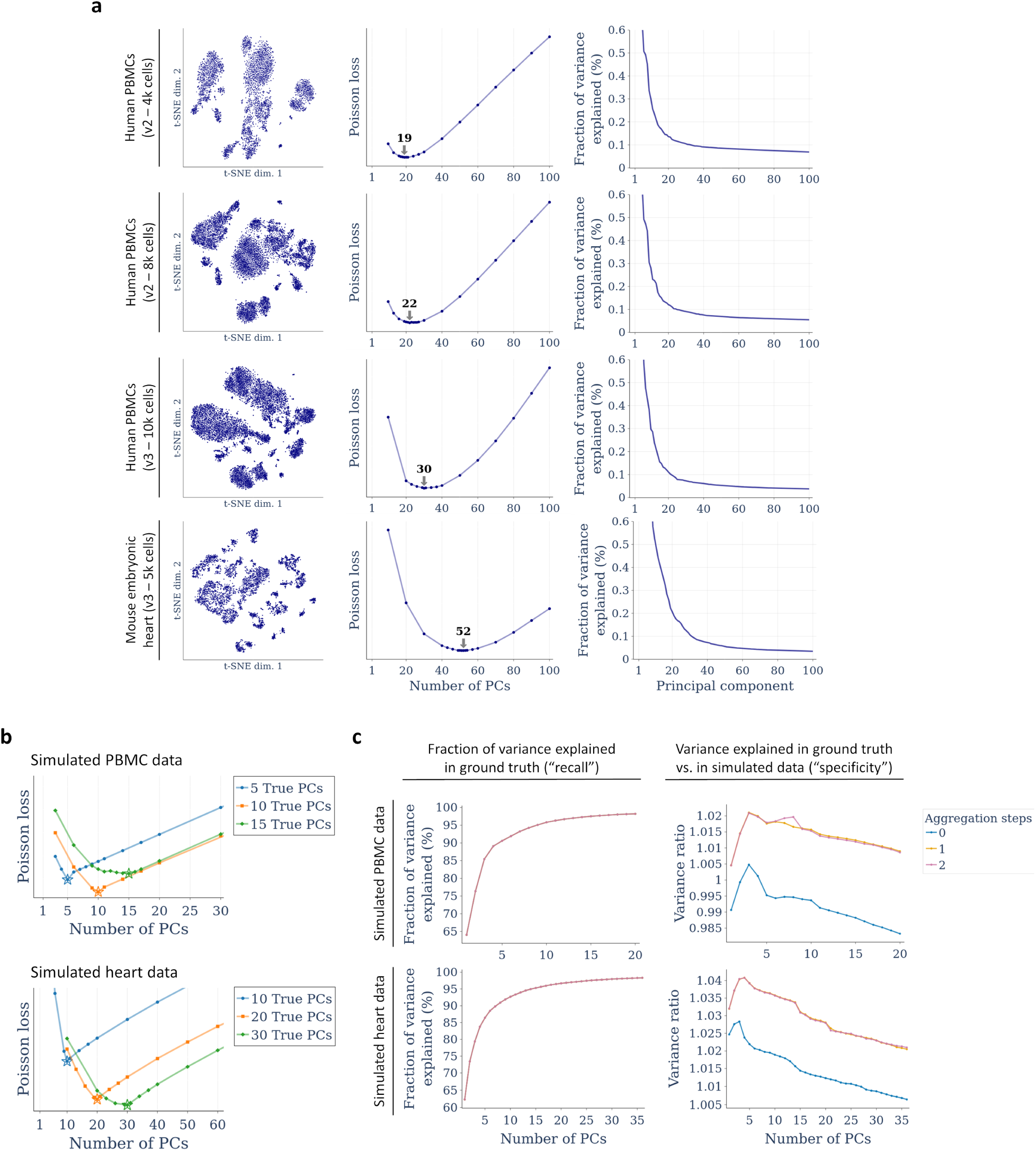
Evaluation of algorithms used in Monet’s core analysis framework. **a** Inference of dataset dimensionality for four scRNA-Seq datasets using molecular cross-validation with a Poisson loss function. Left: t-SNE plots. Center: Poisson loss functions and inferred dimensionality. Right: Percent of variation explained by each PC. **b** Validation of dimensionality inference using simulated PBMC (top) and heart (bottom) datasets with truncated dimensionality. Stars indicate the inferred dimensionality. **c** Quantification of the effect of nearest-neighbor aggregation on PCA performance, using simulated PBMC and heart datasets.

To quantitatively validate the MCV-based inference of dimensionality, I modified a previously described PCA-based approach to simulate scRNA-Seq data using real datasets as templates^3^, which allowed me to generate artificial scRNA-Seq datasets with a truncated and thus clearly defined dimensionality. I simulated human PBMC datasets with dimensionalities of 5-15, and mouse embryonic heart datasets with dimensionalities of 10-30. In all cases, Monet was able to infer the correct dimensionality for the simulated datasets (**Figure 2b**). It should be noted that in real-world datasets where the dimensionality is not artificially truncated, there is typically no sharp transition from dimensions that capture biological expression differences to those that only capture technical noise. This is why the Poisson loss curves look flatter for the real datasets than in the simulation study. Nevertheless, these results showed that Monet’s MCV-based inference of dimensionality provided a valid and more systematic way of determining the dimensionality than the commonly used method of making a guess based on an “elbow plot” that shows the explained variance per PC.

### Monet reduces PCA overfitting by performing nearest-neighbor aggregation

PCA is a highly effective tool for reducing the dimensionality of scRNA-Seq data. In doing so, it separates biological expression differences, captured by the first few PCs, from technical noise, which is captured by higher PCs. In addition to reducing the dimensionality, PCA therefore also denoises scRNA-Seq data^3^. However, when applied to raw UMI counts, higher PCs tend to capture a small fraction of technical noise, which does not exhibit significant correlation structure. This effect can be described as overfitting, as it represents an example of a model capturing unwanted sources of variation. To reduce overfitting, Monet implements a nearest-neighbor aggregation step, which reduces overall noise levels, and thus reduces the extent to which individual PCs capture noise. I performed simulation studies to quantify the extent of this effect in datasets generated with the Chromium v3 technology, and found that the improvements appear relatively minor (**Figure 2c**). The PCs obtained after the aggregation step did not change the percentage of variance explained in the (noiseless) ground truth, however they did lower the percentage of variance explained in the (noisy) simulated data by 2-3%, indicating that overfitting was reduced. A second round of nearest-neighbor aggregation did not improve the results further. Since the simulation method itself relied on PCA, these results are likely biased, and I expect the benefits in real-world applications to be somewhat larger. Additional simulation studies will be required to better quantify this effect.

### Monet enables batch correction by identifying mutual nearest neighbors

As discussed, the analysis of individual scRNA-Seq datasets presents a number of statistical and computational challenges, and there is still surprisingly little consensus as to how to perform even basic tasks such as clustering^4^. However, most single-cell studies require a joint analysis of multiple datasets, for example to compare between different individuals, drug treatments, or genetic backgrounds. In addition, studies often stand to benefit from direct comparisons with previously published scRNA-Seq datasets. In all of these instances, researchers need to adopt strategies to overcome batch effects, which is a catch-all term for all sources of variation that represent technical artifacts rather than true biological expression differences. For example, strong batch effects can be expected when comparing datasets generated using different scRNA-Seq technologies. However, as the precise extent and nature of these sources of variation is typically unknown, batch correction methods generally have to rely on certain assumptions in discriminating technical from biological effects. The development and benchmarking of batch correction methods for scRNA-Seq data is an active area of investigation^16^.

A straightforward and useful method for batch correction was described by Haghverdi et al.^12^, who proposed to identify of pairs of cells from two datasets that represent mutual nearest neighbors (MNNs). These cell pairs would allow the calculation of batch correction vectors, which represent the the batch effect present in a target dataset, relative to a reference. The authors reasoned that after subtracting these batch correction vectors from the cells in the target dataset, any remaining differences between the datasets would represent true biological differences. In effect, this approach assumes that some cell populations are shared between the two datasets, whereas others are unique to either the reference or the target dataset. After applying the batch correction, it should be possible to identify the populations only present in one dataset, but not the other.

Monet implements a modified version of this approach using the *correct_mnn()* function, where the batch correction is performed in PC space, whereas the originally proposed method operates directly on gene expression values. To test Monet’s implementation, I applied batch correction using a human PBMC dataset obtained using the Chromium v3 technology as the reference, and another human PBMC dataset using the Chromium v2 dataset as the target. The difference in technologies results in a strong batch effect, and a clear visual separation of clusters by dataset (**Figure 3a**, top). After applying batch correction, cells can be seen to cluster by cell type, with all clusters containing cells from both datasets (**Figure 3a**, bottom). An advantage of the MNN-based approach to batch correction is that it does not assume that all cell populations are present in both datasets. To demonstrate this on an extreme example, I computationally removed all T cells from either the reference or the target dataset, and applied batch correction again. In both cases, the results for the other cell types were unaffected (**Figure 3b**), confirming that this approach is robust to differences in cell type composition between samples. The batch correction algorithm for these datasets consisting of almost 20,000 cells took approximately 47 seconds. In summary, Monet implements an effective and efficient MNN-based algorithm for batch correction in PC space.

**Figure 3:**
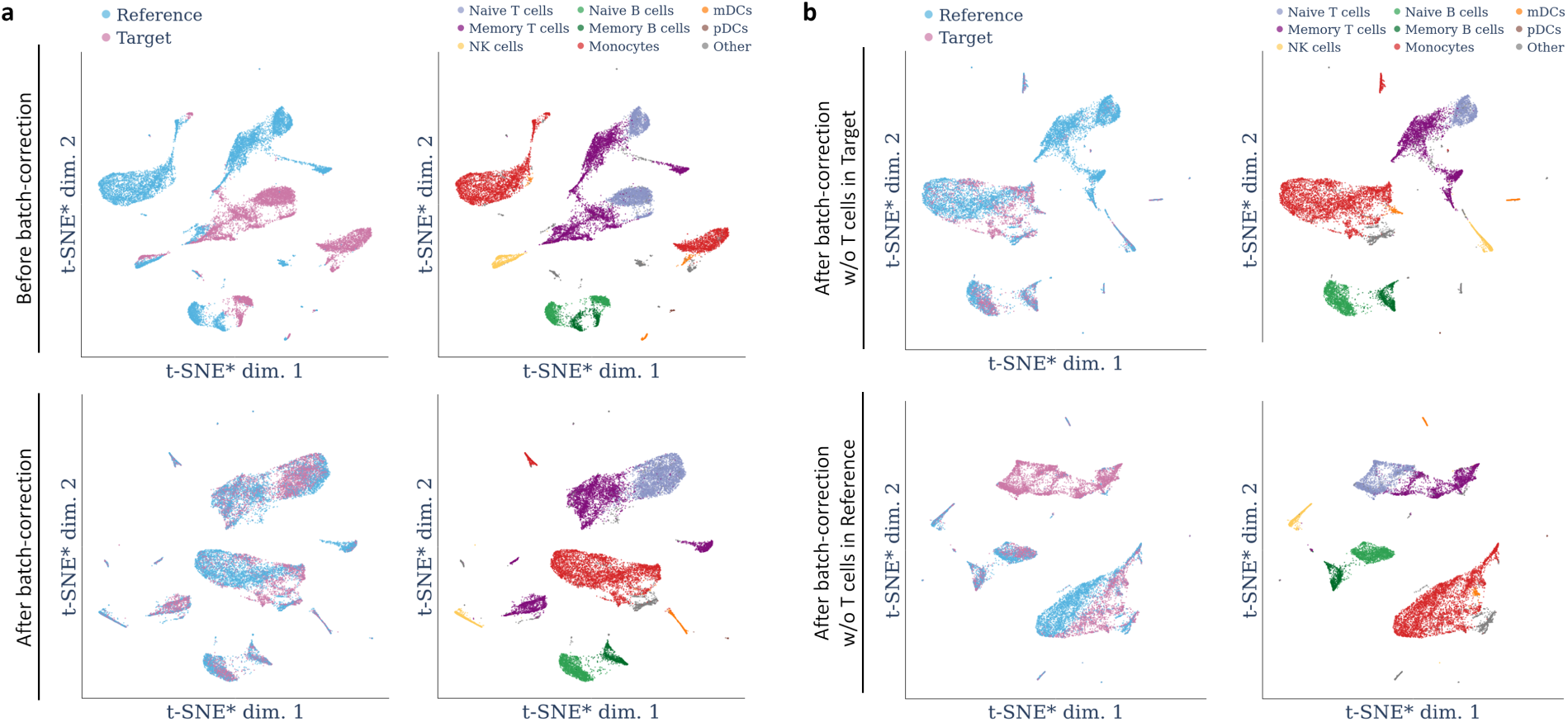
Batch-correction of human PBMC data using mutual nearest-neighbors. The reference and target datasets were obtained using the Chromium v3 and v2 technologies, respectively. **a** t-SNE plots showing data without (top) and with batch-correction of the target dataset. **b** t-SNE plots showing batch correction results obtained after excluding all T cells from either the target (top) or the reference (bottom) dataset. t-SNE*: Exaggerated t-SNE (see Methods).

### Monet enables accurate label transfer between samples from the same tissue

A more supervised approach to overcoming batch effects is to perform clustering on a reference dataset, and to then use machine learning methods to directly transfer cluster labels to other datasets representing samples from the same tissue. The development of such label transfer methods is also a highly active area of investigation^4,17,18^. It is also an area where multiple deep learning-based approaches have been proposed^19,20^. However, as is the case for many scRNA-Seq analysis tasks, it is not clear how much methodological complexity is truly necessary to address this problem, especially in the commonly encountered scenario where a researcher simply wishes to transfer labels between samples from the same tissue.

Monet implements a label transfer method that relies on training a standard k-nearest neighbor (kNN) classifier on the reference data, after projecting it into the latent space represented by a Monet model. A target dataset can then be projected into the same latent space, and labeled using the kNN classifier (**Figure 4a**). Monet uses a default value of *K*=20 for classification, which can be changed by the user. To test this approach, I used the same human PBMC reference dataset as in the batch correction example (see above), obtained using the Chromium v3 technology. I fitted a Monet model and performed clustering, identifying all major cell types present (**Figure 4b**). I then applied the label transfer method to two other human PBMC datasets. First, a dataset obtained using the same Chromium v3 technology, and second, the Chromium v2 dataset that was shown in **Figure 3a** to exhibit strong batch effects. For both datasets, I compared the label transfer results to results obtained from manual clustering (**Figure 4c**). In both cases, Monet correctly identified the vast majority of cells from each cell type (**Figure 4d**), demonstrating that the simple combination of a PCA-based latent space and a kNN classifier was largely successful in transferring cell type annotations, even in the presence of strong batch effects. The most notable exception was the failure to correctly identify approximately 12% of monocytes in the v2 dataset, suggesting that the batch effect for this cell type was too large to allow a reliable classification. It is possible that this could be ameliorated by performing a MNN-based batch correction step (see above) before applying the label transfer method. In future work, I aim to test if this approach indeed leads better results when strong batch effects are present. If so, I plan to make this option directly available in Monet’s label transfer function. It should be noted that the transferred labels appeared to provide a higher cell type resolution than what could be inferred from the t-SNE plot, suggesting that label transfer can help to improve cell type resolution. In summary, the kNN classification-based approach to label transfer implemented in Monet represents a simple and effective tool for transferring annotations between datasets from the same tissue.

**Figure 4:**
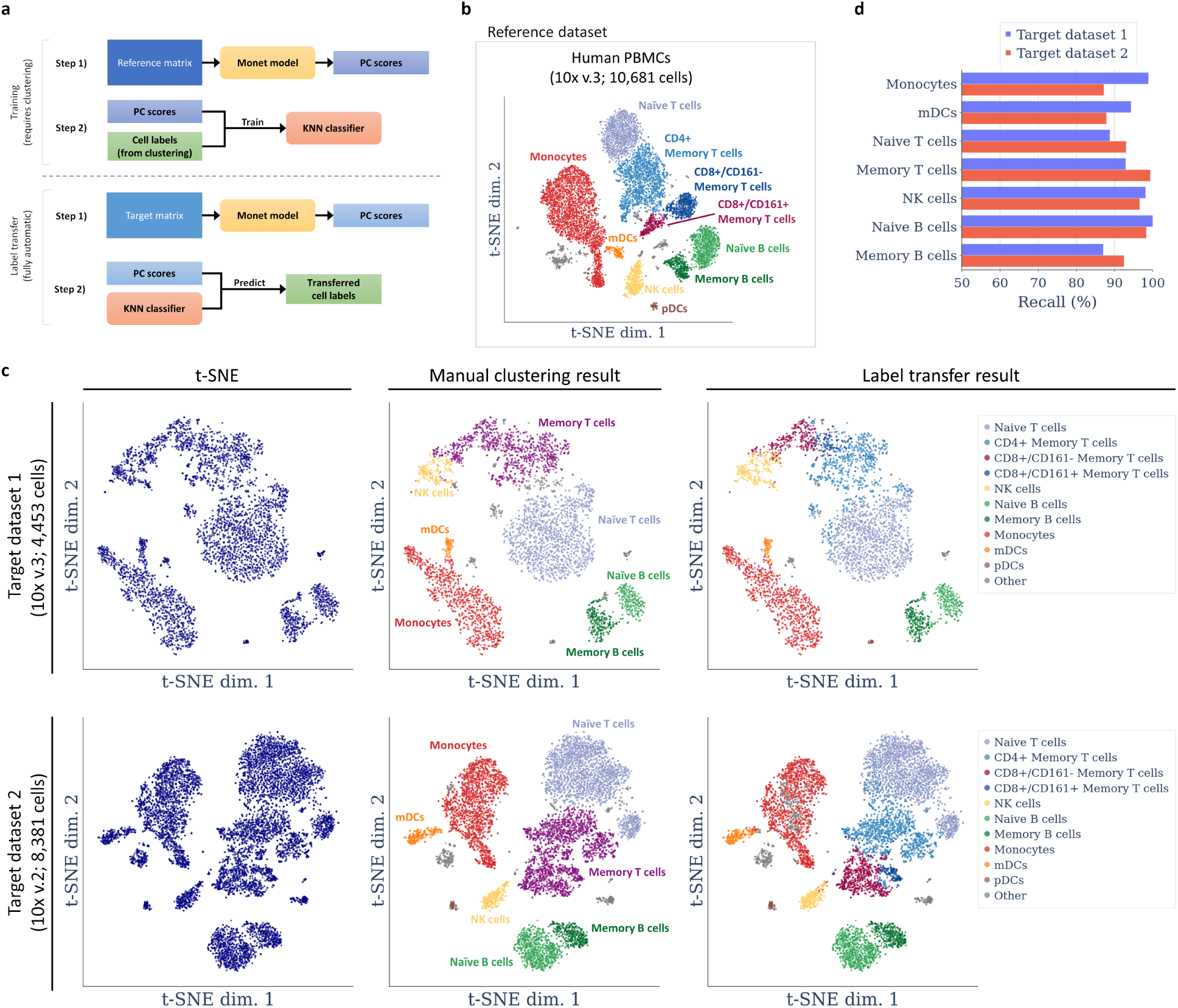
Label transfer using k-nearest neighbor classification, applied to human PBMC data. **a** Schematic of the label transfer approach. **b** Clustering result for the human PBMC reference dataset. **c** Comparison of manual clustering and label transfer results for two human PBMC target datasets. Target dataset 2 (bottom) was obtained using a different scRNA-Seq technology compared to the reference. **d** Quantification of recall per cell type for both target datasets, obtained by treating the manual clustering results as the ground truth.

## Discussion

I have described some of the design choices and features of Monet, an open-source Python package for analyzing and integrating scRNA-Seq data. Most of the analysis tasks currently supported rely on previously described methods, including visualization, clustering, denoising, and batch correction. Therefore, the primary contribution of work does not lie in the development of a novel method for solving a particular analysis task. Rather, this work has focused on bringing together various approaches within the context of a common analysis framework, and on describing a software package that provides concrete implementations of those approaches. The design of Monet was also guided by the idea that analysis methods should not only be effective in accomplishing a particular task, but also computationally efficient (i.e., fast and not too memory-intensive), and not unnecessarily complex. It is my experience that since most analyses are exploratory in nature, algorithms that take several minutes or even hours to finish tend to disrupt the researcher’s analysis workflow. Moreover, the more complex a method is, the more difficult it is to interpret its output and to understand the relationship between raw data and analysis result, which also makes it harder to communicate research results in a transparent fashion. In contrast, methods that rely on simple data transformations and standard machine learning algorithms can be quite easy to understand, at least at an intuitive level. In summary, a fast and simple method is generally preferable to a slow and complex method, especially when it is not clear whether there is a significant difference in accuracy or effectiveness between those methods.

As outlined in the Introduction, “comprehensive” software packages that provide solutions to range of scRNA-Seq analysis tasks play a crucial role in curating and synthesizing methodological and algorithmic knowledge, and in making those methods and algorithms available to the broad community of researchers that employ scRNA-Seq technologies. The popularity of comprehensive analysis packages like Seurat, Scanpy and Monocle means that analysis frameworks and methods implemented by those packages enjoy a far broader visibility and adoption than those only implemented by more specialized packages. However, the number of successful comprehensive scRNA-Seq analysis packages is fairly small, particularly for Python users, resulting in limited diversity within this space. Monet provides a core analysis framework and set of methods that is largely distinct from those implemented by Seurat, Scanpy, and Monocle, and I therefore hope that it contributes to an increase in diversity while providing useful solutions to a number of commonly encountered analysis tasks.

While this work has focused on introducing and evaluating Monet’s core analysis framework and demonstrating its ability to support various analysis tasks, the overall usefulness of the package will also depend on the availability of documentation, tutorials, and software updates. Future work will focus on developing those materials and on maintaining the Monet package.

## Methods

### Overview of the core analysis framework

Several key aspects of Monet’s core analysis framework have been described previously^3,21^. In particular, these previous studies have described the idea of applying PCA after 1) scaling the expression profile of each cell to the median transcript (UMI) count *C* across all cells (“median scaling”) and 2) transforming the scaled values using the *Freeman-Tukey transform*, 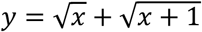 (see below). In the training of a Monet model, this approach is complemented with two additional steps. First, the dimensionality *D* of the data is inferred using molecular cross-validation^9^ (see below). Second, a nearest-neighbor aggregation step is performed to reduce overfitting^3^. Briefly, the PC scores of each cell (of the first *D* PCs) are used to determine the *K* nearest neighbors of each cell, where *K* is automatically adjusted based on the number of cells contained in the dataset and their median transcript count. An “aggregated dataset” with the same number of cells as in the raw data is then obtained, in which each cell expression profile represents an aggregate of *K* cell expression profiles in the raw data. The profile of the *i*’th cell in the aggregated dataset is obtained by aggregating the raw transcript (UMI) counts of the *i*’th cell in the raw data with those of its *K* neighbors. Finally, the aggregated expression profiles are scaled down again to the median transcript count *C* of the raw data. The Monet model then consists of the first *D* PCs of this aggregated and scaled dataset, as well as the transcript count *C*. More specifically, let ***A*** be an *n*-by-*p* matrix containing the aggregated, scaled, and transformed expression values for *n* cells and *p* genes from the training (or reference) dataset. In the singular value decomposition of ***A*, A** = **UΣW**^**T**^, the first *D* PCs are defined as the first *D* columns of ***W***, denoted here as ***W***_*D*_.

To project any scRNA-Seq dataset (typically one from the same tissue as the one used for training the model) into the latent space represented by a Monet model, all cell expression profiles are 1) scaled to the transcript count *C*, 2) transformed using the Freeman-Tukey transform, and 3) projected onto the PCs of the Monet model. More specifically, let ***Y*** by an *m*-by-*p* matrix containing the scaled and transformed expression values for *m* cells and *p* genes from the dataset to be projected. It is assumed here that the set of genes is identical to the set of genes in the training dataset. If necessary, this assumption can be satisfied by removing any unknown genes from the dataset and inserting zero measurements for any missing genes. The PC scores ***S*** for ***Y*** can then be obtained as **S** = **YW**_*D*_.

### Effect of the Freeman-Tukey transform in comparison to the log and Anscombe transforms

The effect and usefulness of the Freeman-Tukey transform on scRNA-Seq data can be appreciated by realizing that scRNA-Seq measurements are associated with significant amounts of technical noise, the exact magnitude of which is expression level-dependent. In order to describe exactly how much noise we expect to observe for a given gene in a given cell, it is helpful to think of scRNA-Seq measurements as random variables, which have specific probability distributions. Based on experiments designed to directly study the noise profile of scRNA-Seq measurements by analyzing “cells” containing identical pools of mRNA (e.g., purified mRNA diluted into droplets), we know that UMI counts can be modeled using the Poisson distribution^1,13,22–24^. Specifically, let *X* represent the UMI count of a particular gene in a particular cell. Then *X* is a Poisson-distributed random variable whose expected value *λ* corresponds to the true (relative) expression level of this specific gene in this particular cell^1,13,23^. The noise level of *X* is described by its coefficient of variation (CV), which due to the Poisson-distributed nature of *X* is inversely proportional to the square root of 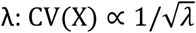. The signal-to-noise ratio is the reciprocal of the coefficient of variation, and therefore directly proportional to the square root of 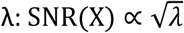. The Freeman-Tukey transform, defined as 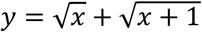, is a variance-stabilizing transform for Poisson-distributed data^14^. Since it is a square root-based transform, it now becomes clear that it approximately weighs each gene expression measurement according to its signal-to-noise ratio. In other words, measurements of highly expressed genes, which are relatively accurate, are given more weight than measurements of lowly expressed genes, which mostly represent technical noise and contain very little information about true expression differences. In fact, after the Freeman-Tukey transform, measurements from lowly expressed genes only have a minimal impact (quantified as the overall proportion of variance associated with those genes). In contrast, after log transform, *y* = ln *x* + 1, those measurements can contribute more heavily than they should based on their signal-to-noise ratio^13^. Virtually all scRNA-Seq analysis workflows that use the log transform rely on a separate step in which the *G* most “informative” genes from the data are selected^2^, and the fact that the log transform tends to assign too much weight to lowly expressed genes appears to be one of the reasons why such a gene selection step is necessary or recommended. The Freeman-Tukey transform, used in combination with median scaling, makes a gene selection step unnecessary. Another transform that accomplishes the same goal is the *Anscombe transform*, 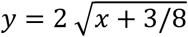. However, this transform has a more extreme effect on the true expression differences (see below).

Aside from weighing gene expression values based on their signal-to-noise ratio, the Freeman-Tukey transform of course also affects the extent to which true expression differences contribute to the analysis. Here, square root-based transforms also differ fundamentally from log-based transforms. The log function grows much more slowly than the square root function, so expression differences in highly expressed genes appear much larger after square root than after log transform, whereas true expression differences in lowly expressed genes can be almost completely lost after square root-based transforms. However, the Freeman-Tukey transform represents somewhat of a compromise between the log-transform and the Anscombe transform, which assigns even more weight to expression differences between highly expressed genes than the FT transform. It should be noted that in the untransformed data, biological variation can be dominated by only a handful of very highly expressed genes.

### Inference of dataset dimensionality using molecular cross-validation (MCV)

Monet’s implementation of MCV closely follows the description provided by Batson et al.^9^, and the reader may refer to their study for a detailed description of this method. Briefly, given a real scRNA-Seq dataset ***X*** with an unknown underlying ground truth of ***X***^*deep*^, the authors describe how to create training and test datasets ***X****’* and ***X****’’* by carefully sampling from the Binomial distribution. In effect, this sampling procedure partitions the observed molecules (UMIs) in ***X*** between ***X’*** and ***X’’***, while allowing for a small overlap. The authors show that if done correctly, ***X****’* and ***X****’*’ represent statistically independent samples of ***X***^*deep*^, mimicking a theoretical scenario in which researchers had performed two independent scRNA-Seq experiments on the same sample (which of course is not possible, as the scRNA-Seq destroys the being cells analyzed).

Once training and a test datasets have been generated, a model of the data (in this case, a PCA-based model) can trained on the training data, and its accuracy can be quantified by calculating an “MCV loss” on the test data. When comparing different models, or models trained using different parameter choices, the most accurate model is that which achieves the lowest MCV loss value. Batson et al. discuss the use of two different loss functions, MSE Loss and Poisson Loss. Monet uses the Poisson Loss function, since each scRNA-Seq measurement can be modeled as a Poisson-distributed random variable (see above). Briefly, assume that the training dataset ***X****’* was used to obtain a *D*-dimensional PCA model of the data, resulting in the PC coefficients ***W***_*D*_ and the corresponding PC scores ***S*** (see above). Let *C*^*test*^ be the median transcript count of the test dataset ***X****’’*. Then let ***M*** be the expression data reconstructed from the scores and coefficients of the first *D* PCs^3^ by 1) reversing the projection onto the first *D* PCs, 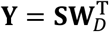, 2) setting all values in ***Y*** below 1 to 1, 2) applying the inverse Freeman-Tukey transform, *x* = (*y*^2^ − 1)^2^/(4*y*^2^), 3) scaling all cell expression profiles to the transcript count *C*^*test*^, and 4) setting all values below 0.001 to 0.001 (this last step avoids taking the logarithm of zero in the loss function, see below). Let ***K*** be the expression data from the test dataset ***X****’’*, also after scaling to the transcript count *C*^*test*^. Then Monet calculates the Poisson loss *L* as follows^9^:

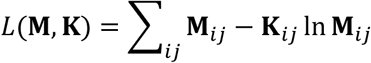

In addition to the choice of loss function, the MCV procedure depends on two parameters. The first parameter is the *capture efficiency p*, which describes what fraction of true mRNA molecules in ***X***^*deep*^ are observed in ***X***. This determines the overlap in molecules between X’ and X’’ that is required to make those samples statistically independent. Batson et al. note that since this overlap is often very small anyway, one could simply choose not to allow any overlap, in which case *p* is ignored. For Monet, I chose a default value of *p*=0.05. I observed that increasing *p* to 0.10, which results in a larger overlap, sometimes resulted in an increased dimensionality, which could either represent a more accurate estimate, or an overestimate due to artificial correlations introduced between the training and test datasets. Since the true capture efficiency of a dataset is generally unknown, *p*=0.05 seemed like a reasonable compromise between not allowing any overlap and running the risk of introducing too much overlap. The second parameter is the *validation ratio α*, which determines the ratio between the proportion of molecules from ***X*** included in the and training datasets, respectively. It is not clear what an optimal choice for *α* is, or if such an optimum exists independently of the dataset being analyzed. However, it is clear that if *α* is chosen too small (e.g., *α*=0.01), then there is a clear risk that the MCV results will become unstable, as so little information is used in calculating the MCV loss. For Monet, I decided to follow Batson et al. and set a default value of *α*=1/9, which mimics a common strategy used in conventional cross-validation that relies on 90%/10% training/test splits of the data. It might be interesting to systematically explore other settings of *α*, for example *α*=0.5.

To infer the dimensionality of a given dataset using MCV, Monet performs a grid search to find the optimum number of *D* between 1 and 100. First, Monet creates five different training/test splits of the data using different random number generator seeds. For each split, Monet then tests an array of possible values of *D*, beginning with the values 10, 20, …, 90, 100. The resulting five Poisson loss values for each parameter setting are then averaged. Then, two successively finer grid searches are performed in the same manner. For example, if the lowest average Poisson loss was observed for *D*=20, then another grid search is performed using *D* values of 13, 16, 19, 21, 24, and 27. If the lowest loss was then observed for D=24, a final grid search is performed using D values of 22, 23, 25, and 26. It appears that the function describing the loss as a function of *D* is generally convex, so this strategy reliably finds the optimum value for *D*, which is the dimensionality that Monet then uses to create a latent space for the dataset.

### Datasets and preprocessing pipeline

The datasets used in this study were as follows:

- “8k PBMCs from a Healthy Donor” (v2-PBMC-8k): Human PBMCs, data generated using the 10x Chromium v2 technology, published by 10x Genomics (https://support.10xgenomics.com/single-cell-gene-expression/datasets/2.1.0/pbmc8k).
- “4k PBMCs from a Healthy Donor” (v2-PBMC-8k): Human PBMCs, data generated using the 10x Chromium v2 technology, published by 10x Genomics (https://support.10xgenomics.com/single-cell-gene-expression/datasets/2.1.0/pbmc4k).
- “10k PBMCs from a Healthy Donor (v3 chemistry)” (v3-PBMC-10k): Human PBMCs, data generated using the 10x Chromium v3 technology, published by 10x Genomics (https://support.10xgenomics.com/single-cell-gene-expression/datasets/3.0.0/pbmc_10k_v3).
- “5k Peripheral blood mononuclear cells (PBMCs) from a healthy donor with cell surface proteins (v3 chemistry)” (v3-PBMC-5k): Human PBMCs, data generated using the 10x Chromium v3 technology, published by 10x Genomics (https://support.10xgenomics.com/single-cell-gene-expression/datasets/3.1.0/5k_pbmc_protein_v3).
- “10k Heart Cells from an E18 mouse (v3 chemistry)” (v3-Heart-10k): Mouse embryonic heart cells, data generated using the 10x Chromium v3 technology, published by 10x Genomics (https://support.10xgenomics.com/single-cell-gene-expression/datasets/3.0.0/heart_10k_v3).

To focus on the expression of protein-coding genes and to reduce matrix size by approximately two thirds, the following gene filtering step was performed. For all datasets, the “Feature / cell matrix (filtered)” file was downloaded from the 10x Genomics website. A list of known protein-coding genes was extracted from the human Ensembl genome annotations, release 97 (http://ftp.ensembl.org/pub/release-97/gtf/homo_sapiens/Homo_sapiens.GRCh38.97.gtf.gz). Each dataset was then filtered to only retain those known protein-coding genes, identified by their Ensembl IDs.

Tor remove low-quality cells and to remove gene expression from genes encoded on the mitochondrial genome, the following quality control steps were performed. A list of 13 protein-coding genes located on the mitochondrial genome was obtained by selecting as all protein-coding genes whose names starts with “MT-”. For all datasets generated using the 10x Chromium v3 technology, individual cells were removed if they had fewer than 2,000 measured transcripts (UMIs), or if more than 20% of measured transcripts originated from those 13 mitochondrial genes. All datasets were then filtered to exclude those 13 genes.

### Visualizations and clustering

Visualizations and clustering analyses using t-SNE and DBSCAN were performed as previously described^21^, using 50 principal components a perplexity of 30. Briefly, the Galapagos clustering workflow was applied, consisting of median scaling, application of the Freeman-Tukey transform, and PCA. Cell type annotations were based on cell type-specific marker genes.

### Generation of simulated data

Simulations were performed by applying the ENHANCE denoising algorithm to a real scRNA-Seq dataset, using the result as the ground truth, and then simulating efficiency and sampling noise to obtain the simulated data^3^. This previously described approach was slightly modified. ENHANCE was applied using the dimensionality (number of PCs) inferred by Monet, rather letting ENHANCE infer the dimensionality using its own heuristic. To validate the MCV approach implemented by Monet, the dimensionality of the ground truth was truncated to a specified number of PCs as follows. The ground truth obtained by ENHANCE can be represented by a set of PC coefficients and scores. To obtain simulated data with a clearly defined dimensionality, only the data represented by the first *D*^*sim*^ PCs were used as the ground truth.

### Benchmarking of PCA performance on simulated data

The percent of biological variance recovered (“variance recall”) using the first *d* components of a PCA model was calculated by first calculating the total variance in the ground truth as follows. The dataset was scaled to the transcript count *C* of the PCA model and then transformed using the Freeman-Tukey transform. Then, the sample variance was calculated for each gene. The total variance was calculated as the sum over all gene variances. To calculate the variance explained by the first d PCs, the PCA model was used to project the ground truth into PC space (see above), and then the sample variance of the PC scores for each of the first *d* PCs was calculated. The variance recall was then calculated as the sum of these variance values, divided by the total variance.

The ratio between the variance explained in the ground truth and the simulate data (“variance specificity”) using the first *d* components of a PCA model was calculated by first calculating the variance explained in the ground truth as before, then calculating the variance explained in the simulated data in analogous fashion, and dividing the two numbers.

### Joint visualization of two datasets with exaggerated t-SNE

t-SNE^25^ visualizations containing a large numbers of cells (e.g., 20,000 or more) tend to have a crowded appearance, with reduced distances between clusters representing different cell types. To avoid this effect, I relied on a modification to t-SNE recently described by Kobak and Berens^26^ that results in a less crowded appearance that is more characteristic of UMAP^27^. This modification consists of setting the rarely modified early exaggeration *α* parameter to 4, and stopping the t-SNE algorithm immediately after the early exaggeration phase. This was accomplished using the t-SNE implementation provided by scikit-learn (*manifold.TSNE*), by setting the *n_iter* parameter to 250, the number of iterations performed with early exaggeration. To visualize cells from two datasets using plotly, I avoided the standard approach in which the cells from each dataset are represented using a separate “trace”, which allows them to be plotted with a dataset-specific color. This results in cells from each dataset being drawn sequentially, and can result in the cells from one dataset occluding cells from the other dataset, producing the illusion that certain clusters only contain cells from one dataset. I therefore used a workaround that produces the same effect while allowing cells to be plotted in completely random order.

### Batch correction using mutual nearest neighbors

Batch correction was performed using the mutual nearest-neighbors approach previously described by Haghvedi et al.^12^, with some modifications. Instead of operating on the raw data after selection of highly variable genes, mutual nearest neighbors are identified after projecting the reference and target datasets into the PC space of the specified Monet model. The output is a dataset containing the batch-corrected PC scores of the cells in target dataset. Briefly, for each query cell from the target dataset, the *K*=20 most similar cells (using Euclidean distance) are identified in the reference dataset, and vice versa. Thus, for each cell a “neighborhood” of cells is defined in the other dataset. MNNs are then defined as pairs of cells who are contained in each other’s neighborhoods. Batch correction is then performed for each cell in the target dataset in the following fashion. First, the five most similar cells in the target dataset for which MNN are available are identified. Then, all MNNs belonging to those five cells are identified. For each MNN, a difference vector is computed by subtracting the PC score vector of the cell in the reference dataset from the PC score vector of the cell in the target dataset. A batch correction vector is then defined as the simple average of those difference vectors. (This represents a simplification of the method proposed by Haghverdi et al., which uses a Gaussian kernel to weigh difference vectors based on the similarity of each MNN to the query cell.) This batch correction vector is then added to the PC score vector of the cell in the target dataset. The batch-corrected PC scores from the target dataset can then be merged with the PC scores from the reference dataset, and t-SNE, exaggerated t-SNE (see above) or UMAP can be performed on this merged PC score matrix. The nearest-neighbor search is implemented using the scikit-learn *neighbors.NearestNeigbhors* class and relies on k-d trees.

### Label transfer using k-nearest neighbor classification

Given 1) a reference dataset in which each cell has been assigned a cell type label (for example, by performing clustering and manually annotating each cluster with a cell type based on marker gene expression), 2) a Monet model (trained on this dataset or another dataset from the same tissue), and 3) a target dataset without cell labels, label transfer was implemented as follows. 1) The reference dataset was projected into the PC space defined by the Monet model. 2) A K-nearest neighbor (KNN) classifier with *K*=20 was trained on the PC scores and cell labels. 3) The reference dataset was projected into the PC space defined by the Monet model. 4) The KNN classifier was used to predict the cell types in the target dataset. The implementation used for KNN classification was the *neighbors.KNeighborsClassifier* class from scikit-learn.

### Software and data availability

Monet is open-source software that is available at https://github.com/flo-compbio/monet under an OSI-compliant 3-Clause BSD License. All analyses were conducted using Python 3.8, and Python packages monet 0.2.0, numpy^28^ 1.18.1, pandas^29^ 1.0.3, scipy^30^ 1.4.1, scikit-learn 0.22.1^31^ and plotly 4.2.1.

The preprocessed scRNA-Seq datasets analyzed in this study will be made available at https://github.com/flo-compbio/monet-paper.

